# High-content protein localization screening *in vivo* reveals novel regulators of multiciliated cell development and function

**DOI:** 10.1101/141234

**Authors:** Fan Tu, Jakub Sedzinski, Yun Ma, Edward M. Marcotte, John B. Wallingford

## Abstract

Multiciliated cells (MCCs) drive fluid flow in diverse tubular organs and are essential for development and homeostasis of the vertebrate central nervous system, airway, and reproductive tracts. These cells are characterized by dozens or hundreds of long, motile cilia that beat in a coordinated and polarized manner (Brooks and Wallingford, 2014). In recent years, genomic studies have not only elucidated the transcriptional hierarchy for MCC specification, but also identified myriad new proteins that govern MCC ciliogenesis, cilia beating, or cilia polarization (e.g. (Choksi et al., 2014b; Chung et al., 2014; Hoh et al., 2012; Ma et al., 2014; Treutlein et al., 2014)). Interestingly, this burst of genomic data has also highlighted the obvious importance of the “ignorome,” that large fraction of vertebrate genes that remain only poorly characterized (Pandey et al., 2014). Understanding the function of novel proteins with little prior history of study presents a special challenge, especially when faced with large numbers of such proteins. Here, we explored the MCC ignorome by defining the subcellular localization of 260 poorly defined proteins in vertebrate MCCs *in vivo*. Moreover, functional analyses that arose from results of the screen provide novel insights into the mechanisms by which the actin cytoskeleton simultaneously influences diverse aspects of MCC biology, including basal body docking, and ciliogenesis.

## RESULTS and DISCUSSION

Rfx family transcription factors play essential roles in vertebrate ciliogenesis (Choksi et al., 2014b), and our recent genomic analysis of Rfx2 function in MCCs revealed over 900 direct target genes, including regulators of MCC progenitor cell movement, ciliogenesis, cilia beating and planar polarity (Chung et al., 2014). Among the Rfx2 target genes, we identified hundreds that either remain poorly characterized or were not previously implicated in MCC biology. To make this “omic” dataset more useful for the study of MCC biology, we systematically determined the subcellular localization in MCCs of 260 of these proteins using live imaging *in vivo*.

### High-content screening of protein localization in MCCs *in vivo*

High content screening of protein localization is a powerful approach for linking genomics to cell biology. This approach has most commonly been applied to invertebrate animals or vertebrate cells in culture (Boutros et al., 2015; Zanella et al., 2010), though more recently it has been applied to intact zebrafish (Choksi et al., 2014a; Clark et al., 2011; Trinh le et al., 2011). To systematically explore protein localization specifically in vertebrate MCCs *in vivo*, we turned to *Xenopus* embryos, in which the epidermis develops as a mix of MCCs and mucus-secreting cells similar to the mammalian airway (Hayes et al., 2007; Walentek and Quigley, 2017; Werner and Mitchell, 2011). Molecular mechanisms of MCC development and function are highly conserved between *Xenopus* and mammals, including humans (e.g. (Boon et al., 2014; Wallmeier et al., 2014; Wallmeier et al., 2016; Zariwala et al., 2013)), but MCCs on the *Xenopus* epidermis develop *in vivo* on the external surface of an externally developing animal. Combined with its molecular tractability (Harland and Grainger, 2011; Wallingford et al., 2010), this feature makes the *Xenopus* embryo an outstanding platform for rapid imaging-based analysis of MCCs.

From our Rfx2 genomic dataset (Chung et al., 2014), we chose candidates proteins encoded by genes that a) require Rfx2 for their expression, b) are bound by the Rfx2 transcription factor, and c) are expressed normally in the *Xenopus* mucociliary epithelium. We selected over 350 candidate genes for which clones were available in the human ORFeome (Rual et al., 2005). Sequencing revealed that roughly 30% of these clones were significantly different from the reported sequence, and these clones were removed from the screen. We used GATEWAY cloning to insert open reading frames into vectors containing N-terminal and/or C-terminal fluorescent protein tags as well as an MCC-specific promoter (Figure 1A). We then injected each of the remaining 260 plasmids directly into blastula stage *Xenopus* embryos (Vize et al., 1991) and observed the localization of the fluorescently tagged proteins by confocal microscopy. All plasmids were co-expressed with membrane-BFP to visualize ciliary axonemes, thus ensuring that localization data reported here is for MCCs (not shown).

**Figure 1.**
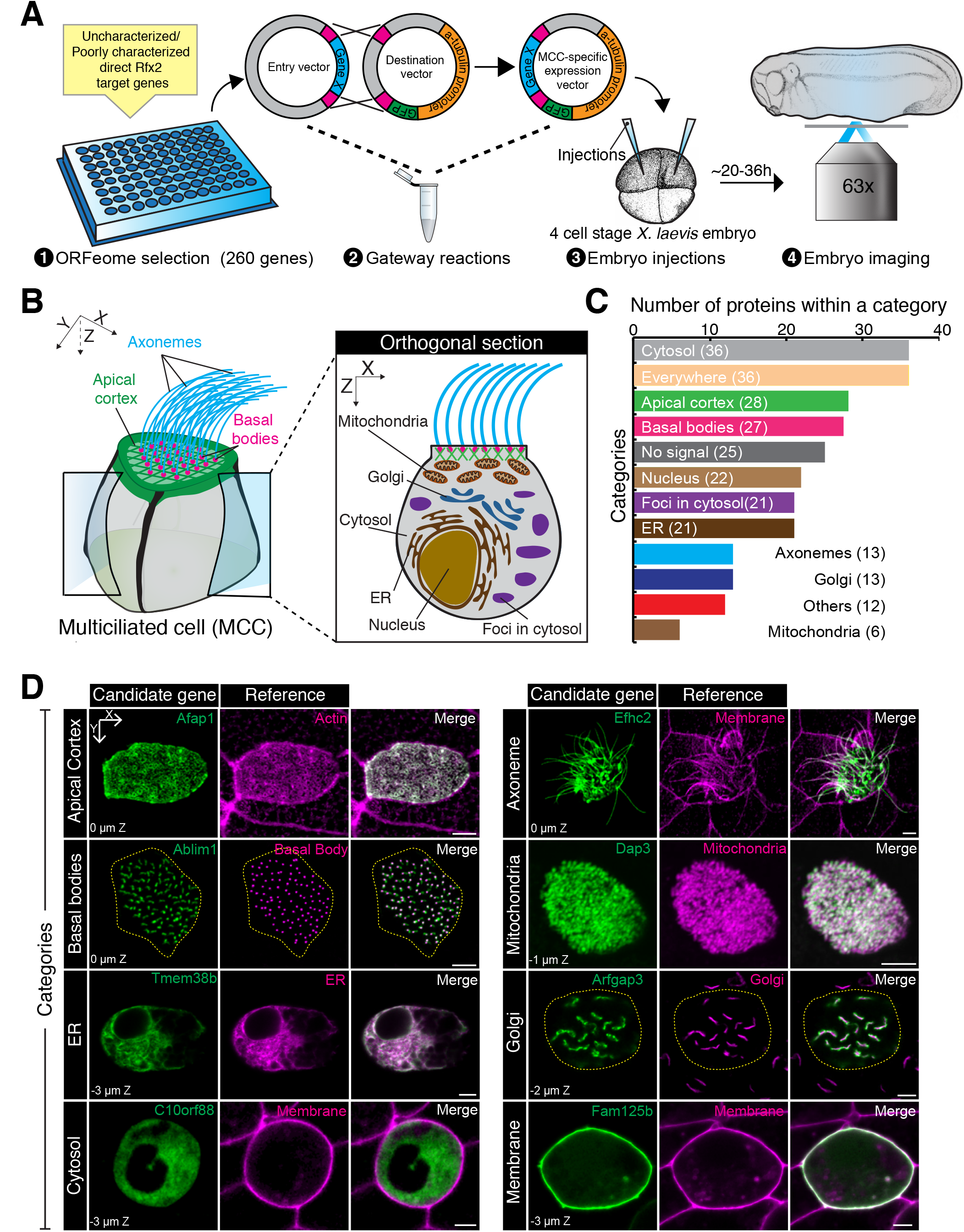
High-content screening of protein localization in MCCs *in vivo*. (**A**) Schematics of the screening pipeline; see Supplemental Methods for details. (**B**) Schematics of MCC subcellular structures, indicating the major distinct subcellular structures identified in our screen. (**C**) Summary of screening results. Out of 260 candidates, 199 showed detectable signal localized to distinct subcellular structures categorized on a histogram. (**D**) Representative localizations of screened Rfx2 targets. Left columns, expression patterns of selected genes; middle columns, expression patterns of reference genes. Afap1 localizes to actin cortex (marked by LifeAct-RFP), Efhc2 to axonemes (marked by CAAX-RFP), Ablim1 to basal bodies (marked by Centrin4-BFP), Dap3 to mitochondria (marked by mito-RFP), Tmem38b to ER (marked by Cal-BFP-KDEL), Arfgap3 to Golgi apparatus (marked by GalT-RFP), C10orf88 to cytosol, and Fam125b to basolateral membrane (marked by CAAX-RFP). Number at the bottom-left corner indicates Z-plane position in reference to the apical domain (0 μm Z). Scale bar, 5μm. Yellow dotted line outlines the cell boundary of a MCC.

Of the 260 candidates tested, 235 were effectively expressed based on GFP or RFP fluorescence, and of these, 199 fusion proteins reported a discrete localization pattern in MCCs (Figure 1B-D; Table S1). While the screen focused on poorly defined and novel proteins, we also included several proteins to serve as positive controls for the specificity of our approach. For example, the axonemal protein Ehfc1 localized to axonemes; the known basal body protein Bbof1/C14orf45 localized to basal bodies; and the transcription factor Dlx3 localized to the nucleus (Table S1)(Bryan and Morasso, 2000; Chien et al., 2013; Ikeda et al., 2005; Zhao et al., 2016).

Consistent with the well-established role for Rfx2 in cilia assembly and function, several proteins localized to the ciliary axoneme and/or the basal body (Fig. 1C, D; Tables S1). For example, Fam166b and Ccdc33 are essentially unstudied and were found strongly restricted to the ciliary axoneme. Mtmr11 and Ankrd45 are similarly uncharacterized and were found specifically at basal bodies. Because the encoding genes are direct Rfx2 targets, these localization patterns strongly suggest a ciliary role, providing the first experimental insights into these proteins’ functions.

In addition to unstudied proteins, some candidates had been previously studied in other contexts, but their linkage here to MCC cilia structure or function may prove informative (Table S1). For example, two known YAP/TAZ-interacting proteins, Ccdc85c and Amotl2 (Wang et al., 2014), were found to localize to basal bodies, potentially shedding new light on the recently described link between YAP/TAZ and ciliogenesis (Grampa et al., 2016; Kim et al., 2015). Likewise, the radial spoke protein Rsph3 was detected in ciliary axonemes as expected, but this protein was also found in MCC nuclei (Table S1), consistent with its reported interaction with Erk2 (Jivan et al., 2009).

Curiously, the majority of candidates tested did not localize to axonemes or basal bodies, but rather labelled diverse cellular compartments, including the cell cortex, the nucleus, the Golgi apparatus, the endoplasmic reticulum, and what appear to be cytoskeletal networks (Figure 1C, D, Figure S1A-D). The preponderance of non-ciliary localization from Rfx2 target genes in MCCs reflects that found previously for FoxJ1 target genes in mono-ciliated cells (Choksi et al., 2014a). Because these organelles are not specific to multiciliated cells, we considered the possibility that these proteins, while encoded by direct Rfx2 target genes, may not actually perform a specialized function in MCCs. However, analysis of *in situ* hybridization data available on *Xenbase* (Karpinka JB et al., 2015) or generated here revealed that many of these genes are in fact strongly and specifically expressed in MCCs (Figure S1E). Moreover, transcription of mRNA for several of the tested candidate genes, including those not localizing to cilia and/or basal bodies, is highly enriched in MCCs of the mammalian airway (Table S2)(Treutlein et al., 2014). Together, these data suggest that even these proteins localizing to common organelles may perform cell-type specific roles in MCCs.

In sum, this screen defines the sub-cellular localization in MCCs of nearly 200 proteins encoded by direct Rfx2 target genes, and thus provides a strong foundation for future studies of MCC biology. As outlined below, these data generate new hypotheses related to diverse human pathologies and also provide new insights into mechanisms of basal body docking and ciliogenesis in MCCs.

### Localization of proteins implicated in human disease

The ciliopathies represent a still-expanding spectrum of human diseases that share an etiology of defective cilia structure or function (Hildebrandt et al., 2011). It is notable then that many of the candidates examined in our screen have been linked to human diseases (Table S3). First, some candidates were implicated in ciliopathies by other studies during the course of our screen. These proteins serve as additional positive controls for our approach and also provide new insights; for example, Armc4 and Ccdc65 are now known as primary ciliary dyskinesia loci, and these proteins are essential for MCC function (Hjeij et al., 2013; Horani et al., 2013). Our data confirm axonemal localization for Armc4 and moreover suggest that the reported cytoplasmic localization of Ccdc65 (Horani et al., 2013) is present in discrete structures, most likely the mitochondria (Table S1). In addition, our screen also examined MCC-specific localization for proteins recently implicated in non-motile ciliopathies. For example, Wdr60 localizes to the base of primary cilia and has been associated with Jeune syndrome and short-rib polydactyly (McInerney-Leo et al., 2013); here we find it similarly localized to the base of multiple motile cilia. Likewise, the transition zone component Tctn3 is implicated in Mohr-Majorski syndrome, and we also find it localized to the base of cilia in MCCs (Table S1). These findings are relevant because even core ciliogenesis machines can also perform MCC specific functions; for example, the IFT protein Wdr35 is required for MCC cilia beating (Li et al., 2015).

Other localization data obtained here suggest new hypotheses related to diverse disease etiologies. Human CERKL mutations are associated with retinitis pigmentosa (Tuson et al., 2004), a known ciliopathy, but the mechanism of CERKL action is poorly defined; our finding here that Cerkl localizes to axonemes and basolateral cell membranes, therefore suggests a novel avenue of inquiry. Likewise, Rfx family transcription factor mutant mice display hearing loss (Elkon et al., 2015), so it is informative that three proteins identified here for their cilia and/or basal body localization are encoded by known human deafness genes (Elmod3, Lrtomta, Cdc14a)(Table S1)(Ahmed et al., 2008; Delmaghani et al., 2016; Jaworek et al., 2013).

Finally, consistent with the overall pattern discussed above, many disease-related proteins encoded by Rfx2 target genes did not have clear localization to cilia or basal bodies. For example, we found that the osteogenesis imperfecta protein Tmem38B (Cabral et al., 2016) localized to the ER in MCCs, which is of potential interest because a recent paper suggests a link between cilia, ER stress, secretory membrane traffic, and low bone density (Symoens et al., 2015; Symoens et al., 2013). Another recent paper proposes an intriguing link between cilia function and scoliosis (Grimes et al., 2016), so it is interesting that two Rfx2 target genes examined in our screen are proposed human scoliosis loci (Karasugi et al., 2009; Kim et al., 2013); Skt localized to the basal bodies and Rims2 to cytoplasmic foci (Table S1).

### Divergent functions for actin regulators in MCCs

MCCs are characterized by a complex apical actin network that is deployed first during assembly of the apical cell surface in nascent cells (Sedzinski et al., 2016; Sedzinski et al., 2017), then for basal body apical migration and docking (Lemullois et al., 1988; Pan et al., 2007; Park et al., 2006; Park et al., 2008), and finally for basal body distribution and planar polarization (Herawati et al., 2016; Turk et al., 2015; Werner et al., 2011). Interestingly, core actin regulators such as the Rho GTPases play distinct roles in these different processes (Pan et al., 2007; Park et al., 2008; Sedzinski et al., 2017). In light of these findings and of the broader link between actin and primary ciliogenesis (e.g. (Avasthi et al., 2014; Kim et al., 2010; Pitaval et al., 2010)), we found it interesting that we identified so many proteins that co-localized to the MCC apical actin network, and that we identified proteins implicated in actin regulation at other sites, such as basal bodies (Table S1). Using these localization patterns as a guide, we explored the mechanisms by which specific proteins direct divergent actin-dependent events during development of MCCs.

### Myo5c controls basal body apical migration

In nascent MCCs, numerous basal bodies are generated in the cytoplasm, and these must migrate apically and dock to the plasma membrane in advance of ciliogenesis (e.g. (Klos Dehring et al., 2013; Sorokin, 1968)). Actin is essential for this apical migration of nascent basal bodies (Antoniades et al., 2014; Boisvieux-Ulrich et al., 1990; Lemullois et al., 1988; Pan et al., 2007; Walentek et al., 2016), but no myosin motor has yet been identified that controls this process. We were interested, then, to identify Myo5c as a basal body localized protein in our screen (Table S1). Curiously, though sequencing of the human Myo5c ORF clone revealed it to be truncated, expression of full-length *Xenopus* Myo5c nonetheless reported a similar localization pattern (Fig. 2A), demonstrating the utility to such high content screening even in light of the known liabilities of large scale clone collections such as the ORFeome.

**Figure 2.**
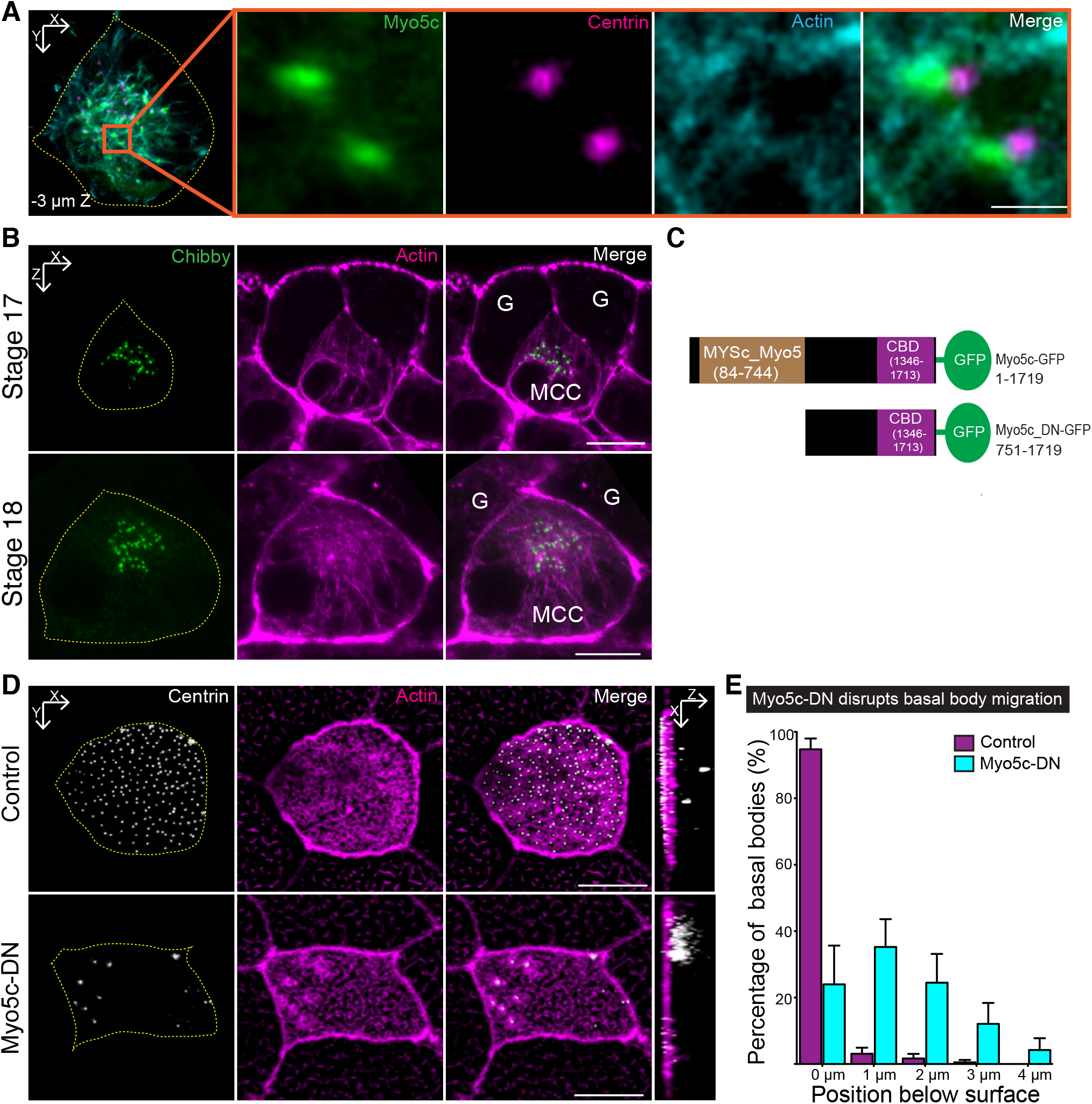
Myo5c localizes to basal bodies and is required for basal body apical migration. (**A**) Myo5c-GFP (green) localized in a proximity to basal bodies (Centrin4-RFP, magenta) and aligned with actin cables (LifeAct-BFP, cyan). Images were taken at stage 19, 3 μm below the apical surface of MCC (outlined by a yellow dotted line). Orange square, zoomed region. Scale bar, 1 μm. (**B**) Sagittal sections of early stage intercalating MCCs showed basal bodies (Chibby-GFP, green) migrating along actin cables (Phalloidin, magenta) to the apical surface of MCC. G, goblet cell. Scale bar, 5μm. (**C**) The dominant negative form of Myo5c was generated by truncating the myosin motor domain of Myo5c (84–744 AA) to disrupt its migration ability but left the cargo-binding domain (1346–1713 AA) intact. (**D**) Overexpression of a dominant negative version of Myo5c (Myo5c-DN-GFP driven by α-tubulin promoter) disrupted basal body migration. In controls(GFP driven by α-tubulin promoter), most of basal bodies (white) docked within the apical actin network (marked by Phalloidin, magenta, and outlined by a yellow dotted line). Upon overexpression of Myo5c-DN, basal bodies failed to migrate apically and accumulated below the apical surface, see orthogonal views. Scale bar, 10 μm. (**E**) Quantification of basal body positions in controls and upon overexpression of a Myo5c-DN in MCCs. More than 75% of the basal bodies in Myo5c-DN overexpressing cells failed to migrate to the apical surface of MCCs (outlined by a dotted yellow line). The average depth of basal bodies increased from 0.08±0.06 μm in controls to 1.37 ±0.35 μm below the apical domain (reference position, 0 μm) in Myo5-DN, (P<0.001; control, n=13 cells; Myo5C-DN, n=14 cells, N>5 embryos. Data represent mean and SD).

Myo5c is a well-studied regulator of epithelial cell biology and though it is expressed in the mammalian airway (Rodriguez and Cheney, 2002), it has never been studied in MCCs. Like other Type V myosins, Myo5c is known to control actin-based movements of intracellular cargoes, including organelles, such as melanosomes and secretory vesicles (Bultema et al., 2014; Marchelletta et al., 2008; Rodriguez and Cheney, 2002; Sladewski et al., 2016). Thus, the observed localization to basal bodies suggested that Myo5c might play a similar role in basal body transport. Consistent with this idea, we observed Myo5c localizing to basal bodies during their apical migration, at stages when basal bodies are associated with actin cables (Figures 2A, B).

A recent report suggests that Class V myosins are required for primary ciliogenesis (Assis et al., 2017), so we sought to test the function of Myo5c in MCCs. Because these myosins play multiple roles in many cells types, we circumvented any potential early effects on development by express a dominant-negative version of Myo5c (Rodriguez and Cheney, 2002) specifically in MCCs using our MCC-specific promoter (Figures 2C,D). We used the cargo-binding Myo5C tail as a dominant-negative (Fig. 2C), as this strategy has been widely deployed for examining unconventional myosin functions (Rodriguez and Cheney, 2002; Rogers et al., 1999; Wu et al., 1998). Strikingly, expression of this construct in MCCs function resulted in profound defects in basal body docking (Figure 2E). This identification of a specific myosin motor involved in actin-based apical migration of basal bodies demonstrates the value of large-scale protein localization screening to identify novel, cell type-specific functions for broadly acting proteins.

### Dennd2b controls axonemogenesis after basal body docking

Following apical surface emergence and basal body docking, MCCs extend their many dozens of axonemes in a process similar to primary ciliogenesis. Interestingly, in many cell types, actin has been suggested to play diverse roles in axonemogenesis (i.e. extension of the axoneme *after* basal body docking)(Avasthi et al., 2014; Kim et al., 2015; Kim et al., 2010; Pitaval et al., 2010), though such functions have not been reported for actin in MCCs. In light of these findings, we explored additional actin co-localizing proteins from our screen. Among these, Dennd2b (previously known as St5) was particularly interesting, as it has been associated with actin-based structures in mesenchyme cells in culture (Ioannou et al., 2015), but it has not been studied in MCCs or any other ciliated cell type.

MCC apical actin is comprised of two elements, an apical meshwork and sub-apical foci (Werner et al., 2011), and we found that full length *Xenopus* Dennd2b localized to both structures (Figure 3A). Knockdown of Dennd2b severely disrupted the assembly of sub-apical actin foci, but elicited only a mild disorganization of the apical actin meshwork (Figures 3B, C, S2A). Interestingly, Dennd2b knockdown also elicited a disruption of basal body planar polarity (Figures 3D, E), an effect that may be direct or may be secondary to a loss of fluid flow, as both apical actin and cilia beating are required for the fluid-flow dependent refinement of basal body planar polarity (Mitchell et al., 2007; Werner et al., 2011).

**Figure 3.**
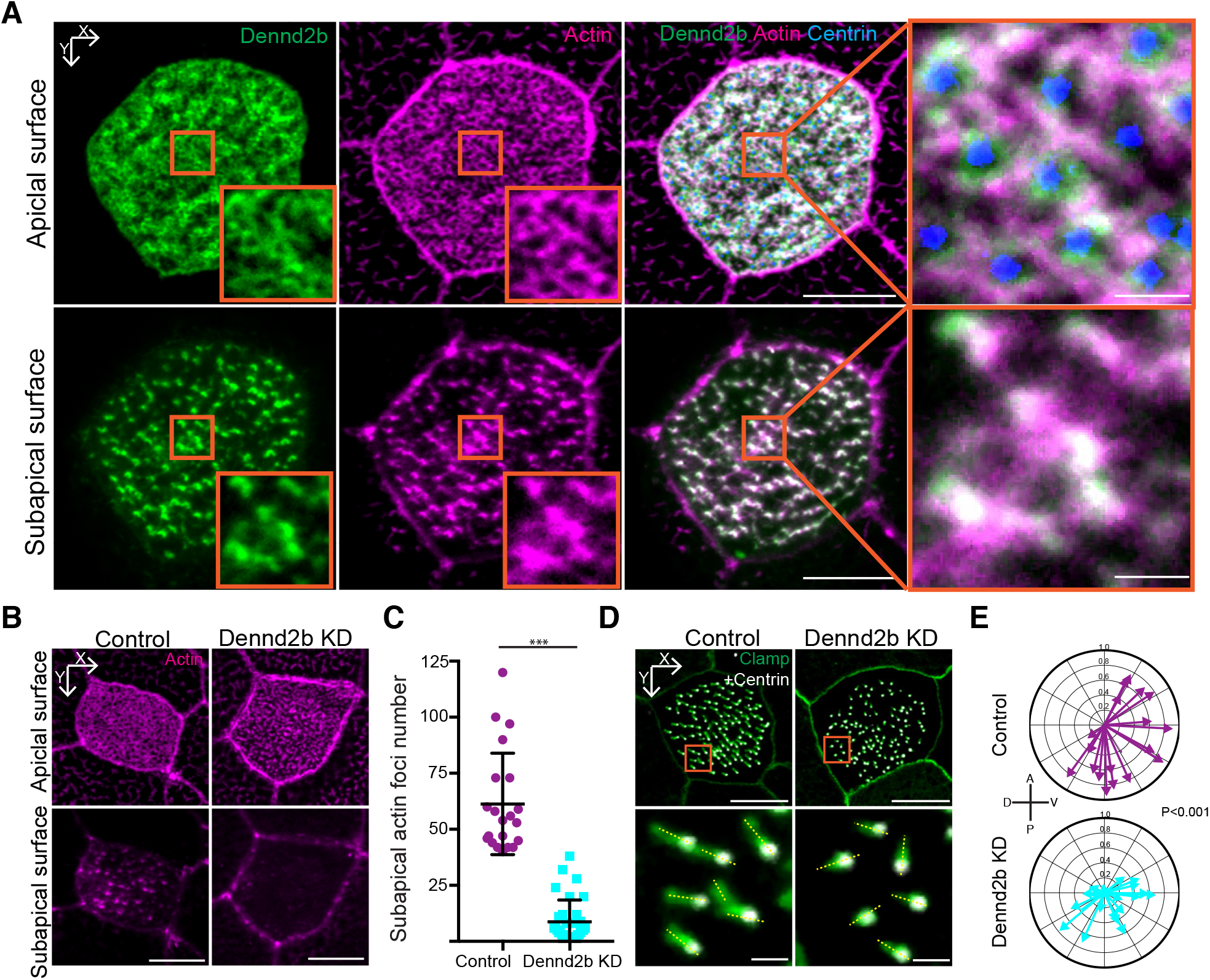
Dennd2b localizes to apical actin networks and is required for basal body planar polarity. (**A**) GFP-Dennd2b (green) localized to apical and subapical actin network (marked by phalloidin, magenta). Scale bar, 10 μm (in zoomed region, scale bar, 1 μm). Images were taken at stage 32. (**B**) Dennd2b KD reduced the number of subapical actin foci (actin marked by phalloidin, magenta). Scale bar, 10 μm. (**C**) The number of subapical actin foci was decreased from 61.3±22.7 in controls to 8.7±9.7 in Dennd2b KD (P<0.001; Control, n=21 cells; Dennd2b KD, n= 33 cells, N>5 embryos). Data represent mean and SD. (**D**) Dennd2b KD disrupts basal body orientation. Orientation of basal bodies was determined by measuring angle (yellow dotted line) between a basal body (marked by Centrin4-RFP, white) and corresponding rootlet (marked by Clamp-GFP, green) in respect to the horizontal line. Scale bar, 10 μm (top panel), 1 μm (bottom panel). (**E**) Quantification of basal bodies orientation. Each arrow represents one cell, where length indicates uniformity of measured angle in that cell (resultant vector). The mean resultant vector value was decreased from 0.71 ± 0.18 in controls to 0.39 ± 0.2 in Dennd2b KD. (P<0.001; Control, n=19 cells; Dennd2b KD, n= 22 cells, N>5 embryos).

Surprisingly, unlike disruption of Myo5C (above), Dennd2b knockdown had no effect on basal body docking (Figure S2D). Rather, disruption of Dennd2b specifically suppressed elongation of ciliary axonemes from docked basal bodies(Figures 4A, B, S2A,C), suggesting that Dennd2b represents a novel actin-related activity that is required for MCC axoneme assembly. Targeting of Dennd2b using CRISPR (e.g. (Tandon et al., 2016)) elicited a similar phenotype (Figures S2B,C).

**Figure 4.**
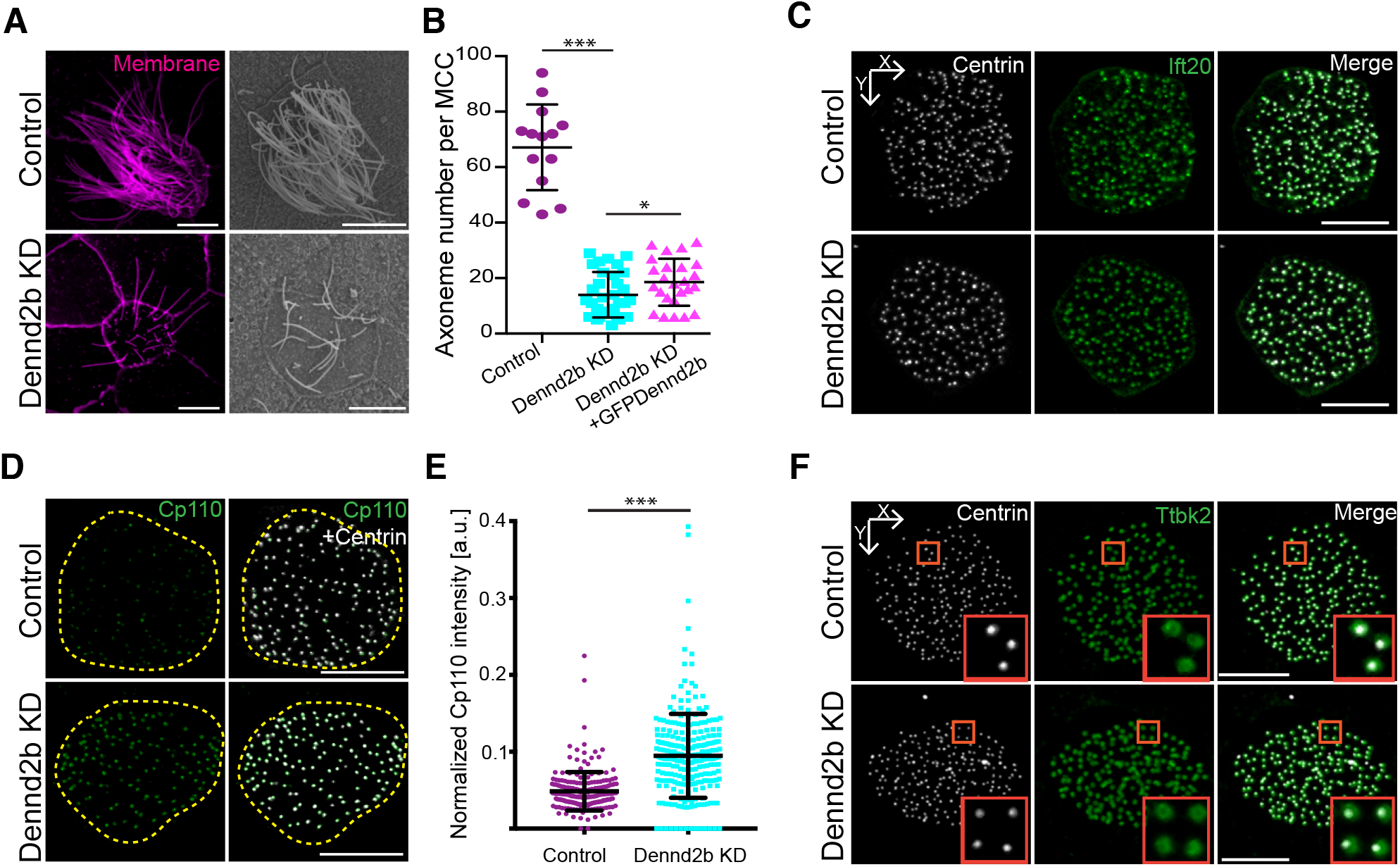
Dennd2b is required for axonemogenesiss and reduction of Cp110. (**A**) Perturbation of Dennd2b disrupted ciliogenesis. Left panel, axonemes visualized by CAAX-RFP, magenta. Right panel, SEM of a control MCC and upon Dennd2b KD. Scale bar, 10 μm. (**B**) The axoneme number per MCC was significantly reduced from 67.1 ± 15.4 in controls to 14.0 ± 8.2 in Dennd2b KD, (P<0.001; Control, n=14 cells; Dennd2b KD, n= 32 cells, N>5 embryos). Number of axonemes was increased by rescue expression of GFP-Dennd2b with a-tubulin promoter to 18.6 ± 8.5. (P<0.05; Dennd2b KD with GFP-Dennd2b, n=25 cells, N>5 embryos). Data represent mean and SD. (**C**) Dennd2b KD did not disrupt the recruitment of Ift20 (Ift20-GFP, green). Scale bar, 10 μm. (**D**) Dennd2b KD increased the level of GFP-Cp110 (green) at basal bodies (marked by Centrin4-RFP, white). Scale bar, 10 μm. (**E**) The normalized Cp110 intensities around basal bodies increased from 0.046 ± 0.045 to 0.093 ± 0.098. (P<0.001; Control, n=1430 intensities from 8 cells; Dennd2b KD, n= 1413 from 10 cells, N>5 embryos). Data represent mean and SD. (**F**) Dennd2b is dispensable for Ttbk2 (GFP-Ttbk2, Green) recruitment at basal bodies.

Among the recent studies implicating actin in axonemogenesis is one showing that actin is required for recruitment of intraflagellar transport (IFT) proteins to the base of cilia in *Chlamydomonas* (Avasthi et al., 2014). However, we observed no reduction in basal body recruitment of the IFT-B subunit Ift20 after Dennd2b knockdown (Figure 4C). Other recent work in both *Xenopus* and mammalian MCCs suggests an interplay between the actin cytoskeleton and the centrosomal protein Cp110 (Cao et al., 2012; Chevalier et al., 2015; Song et al., 2014). Cp110 is an especially attractive candidate because, as tight regulation of Cp110 protein levels was recently found to be critical for MCC ciliogenesis (Walentek et al., 2016).

Strikingly, we found that knockdown of Dennd2b resulted in a significant increase in Cp110 at basal bodies in *Xenopus* MCCs (Figures 4D, E). Cp110 was also recently shown to interact with Fak (Walentek et al., 2016), which is known to link MCC basal bodies to actin (Antoniades et al., 2014), however we found that Fak localization was not disturbed after Dennd2b knockdown (Figure S2E). Likewise, Ttbk2 is known to be required for Cp110 removal (Goetz et al., 2012), but Ttbk2 was also not lost from basal bodies after Dennd2b knockdown(Figure 4F). Thus, our data suggest that Dennd2b is a novel regulator of Cp110 levels at basal bodies, and thereby plays a key role in MCC ciliogenesis.

## CONCLUSION

Here, we used high-content protein localization screening *in vivo* to rapidly advance our understanding of the MCC ignorome. We screened 260 proteins, identifying specific localization patterns for 199, including localization to axonemes, basal bodies, the cell cortex, the cytoskeleton, and specific compartments in the cytoplasm (Figure 1). Directed studies emerging from this screen provide new insights into mechanisms by which the actin cytoskeleton directs MCC development and function (Figures 2–4).

Moreover, because our previous work in *Xenopus* MCCs predicted a broad role for the CPLANE proteins in human ciliopathy (Park et al., 2006; Toriyama et al., 2016), it is notable that two previous studies have implicated Dennd2b/St5 in undefined birth defect syndromes reminiscent of ciliopathies (Gohring et al., 2010; Kleczkowska et al., 1988). Strikingly, one of these patients presented with chronic otitis media and the other with recurrent respiratory infections (Gohring et al., 2010; Kleczkowska et al., 1988); both are hallmarks of motile ciliopathies, so our data here from MCCs may be especially informative.

Finally, given the key role of MCCs in normal airway physiology, our data also provide novel hypotheses for studies of acquired airway diseases. For example, Cp110 expression levels are altered in chronic rhinosinusitis (Lai et al., 2011), so our links to Dennd2b and apical actin may be informative. Moreover, this study is also informative because apical actin is a direct target of pneumoccoal infection in the airway (Fliegauf et al., 2013). In sum, our work highlights the power of combining large-scale image-based screening *in vivo* with more traditional -omics approaches for rapid exploration of the ignorome in cell and developmental biology and provides a rich dataset for further exploration of MCC development and function.

## EXPERIMENTAL PROCEDURES

Please see the Supplemental Experimental Procedures. All animal experiments were approved by the IACUC of the University of Texas at Austin, protocol no. AUP-2012-00156

## AUTHOR CONTRIBUTIONS

F.T, J.S, E.M. and J.B.W. conceived the project and designed the experiments. F.T. performed the screen and most of the experiments. J.S. contributed to experiments and data analysis. Y.M. performed SEM on *Xenopus* embryos. F.T., J.S. analyzed the data. F.T., J.S., and J.B.W. wrote the paper. E.M. provided the ORFeome and commented on the manuscript.

## ACKNOWLEDGMENTS

We gratefully acknowledge members of the Wallingford lab for helpful advice. This work was supported by grants to J.B.W from the NHLBI and NICHD and to E.M.M. from the NIH, NSF, CPRIT and Welch Foundation (F-1515).

